# Revealing variants in SARS-CoV-2 interaction domain of ACE2 and loss of function intolerance through analysis of >200,000 exomes

**DOI:** 10.1101/2020.04.07.030544

**Authors:** Elizabeth T. Cirulli, Stephen Riffle, Alexandre Bolze, Nicole L. Washington

**Affiliations:** Helix, 101 S Ellsworth Ave Suite 350, San Mateo, California 94401

## Abstract

Our researchers took a look at a sequence of DNA known as the *ACE2* gene. This gene is most well known for its role in regulating blood pressure. But in recent times, it’s drawn a lot of attention from the scientific community because it may also serve as a doorway of sorts, enabling viruses like SARS-CoV-2 to infect cells. Our researchers looked at the *ACE2* gene in more than 200,000 people, comparing their exact DNA sequences to see where there are differences among people. Variation in the DNA sequence of a gene is common and is sometimes meaningless. But other times, small changes in the DNA sequence can alter the protein that is made from that gene. In this case the *ACE2* gene makes the ACE2 protein, which is what the SARS-CoV-2 virus interacts with. We found a lot of variation between individuals and checked to see if that variation coincided with any traits (i.e., people with variant X tend to have high blood pressure more often than people without variant X). All of the traits we looked at were non-COVID-19-related traits, meaning we haven’t asked these people anything about COVID-19 yet (this is because these DNA sequences were collected before the pandemic).

We found that there are a number of variations observed among people in a specific part of the *ACE2* gene. These variations are expected to alter the shape or functionality of a specific part of the ACE2 protein: The part that interacts with the SARS-CoV-2 virus. We don’t yet know what the real-life significance of this variation is, but it’s possible that these variants decrease the protein’s ability to interact with the SARS-CoV-2 virus, thus decreasing the person’s likelihood of being infected. We can speculate that there will be a spectrum of vulnerability to COVID-19 among people, where some people are more vulnerable than others, and that variants in this part of the *ACE2* gene may be one of the reasons. The research we presented here shines a light on this part of the *ACE2* gene and may give future researchers a direction to go in as they try to figure out what makes people vulnerable to COVID-19 and similar viruses.

## COVID-19 host genetics

In recent months, our world has been hit by the novel coronavirus SARS-CoV-2, which causes the disease COVID-19. Multiple groups around the world, including Helix, are currently working to genotype and/or sequence individuals who have been infected with SARS-CoV-2, and make their datasets available, in order to study potential host genetics and susceptibility factors for COVID-19 ^1–3^. During the HIV epidemic, researchers were able to discover that approximately 1% of European ancestry individuals were immune to HIV infection due to a deletion in the host coreceptor CCR5^4–6^. Similar genetic findings could help us identify individuals who are not at risk for COVID-19 or who are predisposed to certain disease outcomes. While no one has yet sequenced infected individuals, we are able to examine the naturally occurring variation at the population level in genes suspected to play a role in SARS-CoV-2 infection.

## The importance of *ACE2*

Currently, the human host receptor for this virus is thought to be Angiotensin I converting enzyme 2 (*ACE2*) ^7,8^, a 19-exon gene encoding a protein of 805 amino acids, with only one major protein-coding isoform^9^. ACE2 is a carboxypeptidase transmembrane protein that cleaves angiotensin I and II, controls vasoconstriction and blood pressure (and therefore hypertension), and was also previously established as the host receptor for SARS-CoV (from the 2002 epidemic)^10^. It has high expression levels in multiple tissues, especially those of the small intestine and testis^11^. Multiple common variants in linkage disequilibrium (LD) with each other are already known to be associated with *ACE2* expression levels in the brain, but not in other tissues, especially those with high expression of the gene^12^. Importantly, *ACE2* is on the X chromosome. Because males have only one copy of the X chromosome and females have two, this means that any males carrying one copy of a variant in this gene are more likely to see its effect on their phenotype than would a female carrying one copy of the same variant. Finally, mouse models with a presumed knockout of the *ACE2* gene showed cardiac, metabolic, muscle and pulmonary phenotypes, including pulmonary vascular congestion and increased lung weight, and they were more likely to die early of congestive heart failure after transverse aortic constriction^13^.

## Genetic variation in *ACE2*

At Helix, we queried our database of more than 200,000 exomes of unrelated people to identify the frequencies of different coding variants, including CNVs, in *ACE2*. We have identified 332 variants that affect the coding sequence of this gene, 16 of which are loss of function (LoF), five of which are CNVs, and 174 of which are not found in either gnomAD v2.1.1 (115,000 exomes and 15,000 genomes) or gnomAD v3 (70,000 genomes) (Figure 1) ^14,15^. The frequencies of these variants in unrelated individuals from populations of different ancestries can be found in the Table S1. Importantly, we compared the variants we found to the recently reported structure of ACE2 bound to SARS-CoV-2^16,17^. We identified 11 coding variants in 83 individuals that changed the specific amino acids shown to physically interact with SARS-CoV-2, and an additional 29 variants in 1,885 individuals that were within two amino acids of these crucial sites (Figure 2). The most common among these variants is chrX:15600835:T:C / p.K26R, which has an allele frequency of ~0.5% in multiple populations and changes the amino acid directly adjacent to a SARS-CoV-2-interacting amino acid but is not predicted to be damaging. Of the 40 variants in the binding regions, 13 were predicted to be damaging, and 3 of these were specifically on the interactive amino acids. Notably, one of the damaging variants that directly affects a SARS-CoV-2 interacting amino acid, chrX:15600857:A:G / p.S19P, is very rare or absent in most populations but has an allele frequency of 0.1-0.2% in those with African ancestry. Variants in these interacting regions may change the strength of the bond with SARS-CoV-2, which could have effects ranging from being protective to increasing risk, and they are interesting genomic targets for further study.

**Figure 1.**
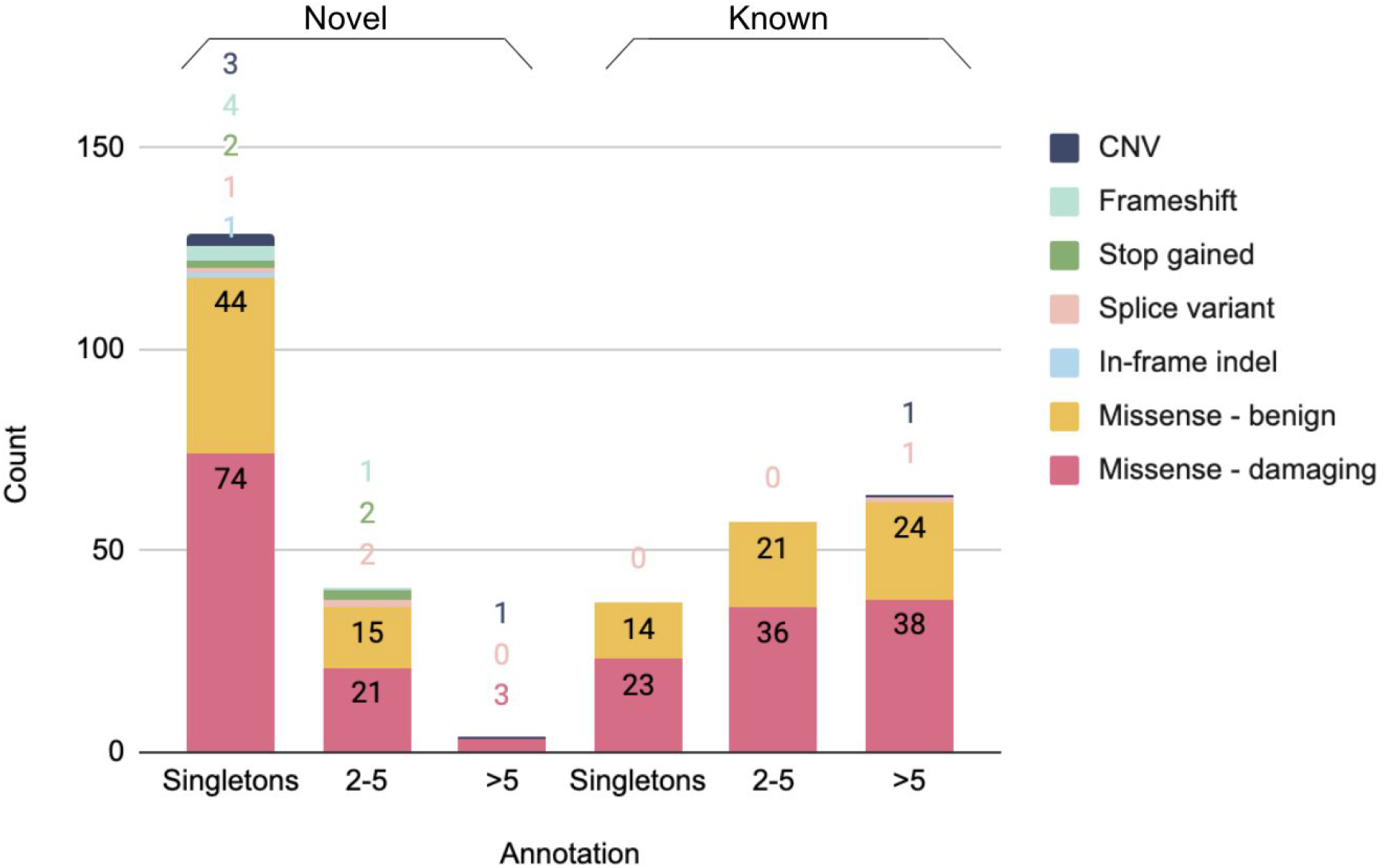
*ACE2* variants found in the Helix dataset of >200,000 unrelated individuals. Count of coding variants by type, grouped by singleton, 2-5 carriers, or >5 carriers and whether novel or in gnomAD database(s).

**Figure 2.**
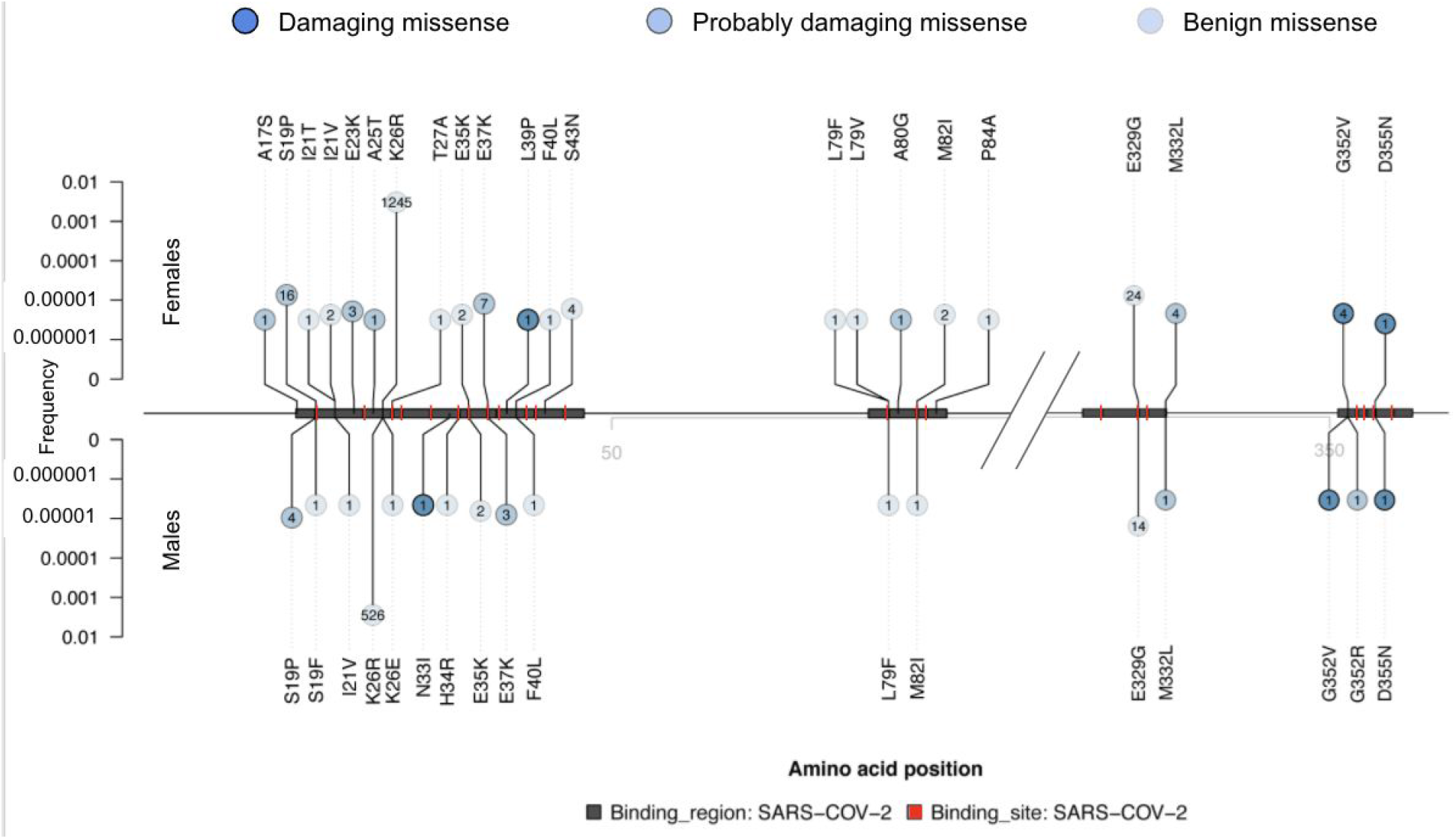
Lolliplot^18^ of missense variants in the regions of ACE2 that bind to SARS-CoV-2. The amino acid positions along ACE2 are shown along the horizontal middle line, SARS-CoV-2 binding regions^16,17^ are shown as gray boxes, and the specific amino acids that bind to SARS-CoV-2 are highlighted in red. For clarity, only the binding regions are depicted (0-95 and 315-360 of 805 amino acids). The vertical line height above the middle line represents allele frequencies in females, and below are allele frequencies in males (log scale). The missense variants in these regions are annotated as damaging (SIFT^19,20^ <0.05 and Polyphen2^21,22^ >0.85; dark blue), probably damaging (SIFT <0.05 or Polyphen2 >0.15; medium blue), or benign (SIFT >0.05 and Polyphen2 <0.15; light blue). All coding variants in the Helix dataset that overlap the binding regions are shown, with case counts of each variant indicated in the bubble, and amino acid change indicated. Y axis is log scale.

## *ACE2* hemizygous LoF depletion

Of these variants, we identified only a single individual with an *ACE2* full-gene deletion. As it turns out, the individual was female, and retained one functioning copy of *ACE2* on their other X chromosome. However, if a male had the gene deleted, then they would be entirely missing this gene. Likewise, other LoF variants would result in no working copies of *ACE2* in a male but might have no effect in a female. Looking at >200,000 individuals sequenced at Helix, we find that predicted LoF variants collectively have an allele frequency of 0.013% in females but 0.007% in males for those of European ancestry, and <0.05% in non-European ancestry females and males (Figure 3). Among high-confidence LoF variants as annotated by LOFTEE^23^, the frequencies were 0.01% in European ancestry females and 0.005% in males. No females were homozygous or compound heterozygous for LoF variants. Additionally, we observe no frameshift variants in males despite finding them in six females (Figure 4); most of the LoF variants seen in males are splice variants, and only a single male was found to have a premature stop codon, in the C-terminal collectrin domain. This pattern is consistent with hemizygous depletion of LoF variants in this gene (Fisher’s exact p=0.002), meaning that the complete loss of this gene is likely deleterious or even lethal. Furthermore, this result is consistent with the data available from gnomAD’s ~185,000 exomes and genomes, where the LoF allele frequency is 0.009% in females of European ancestry as compared to 0.003% in males (non-European ancestries, <0.02% in females and males) and the UK Biobank’s ~50,000 exomes, where it is 0.013% in females of British ancestry and 0.006% in males, and by its published probability of being loss-of-function intolerant (pLI) score of 1^24,25^. Because LoF variants are so rare in this gene, the exact frequency in specific non-European ancestry populations, which have smaller sample sizes available in these large databases, cannot be precisely calculated.

**Figure 3.**
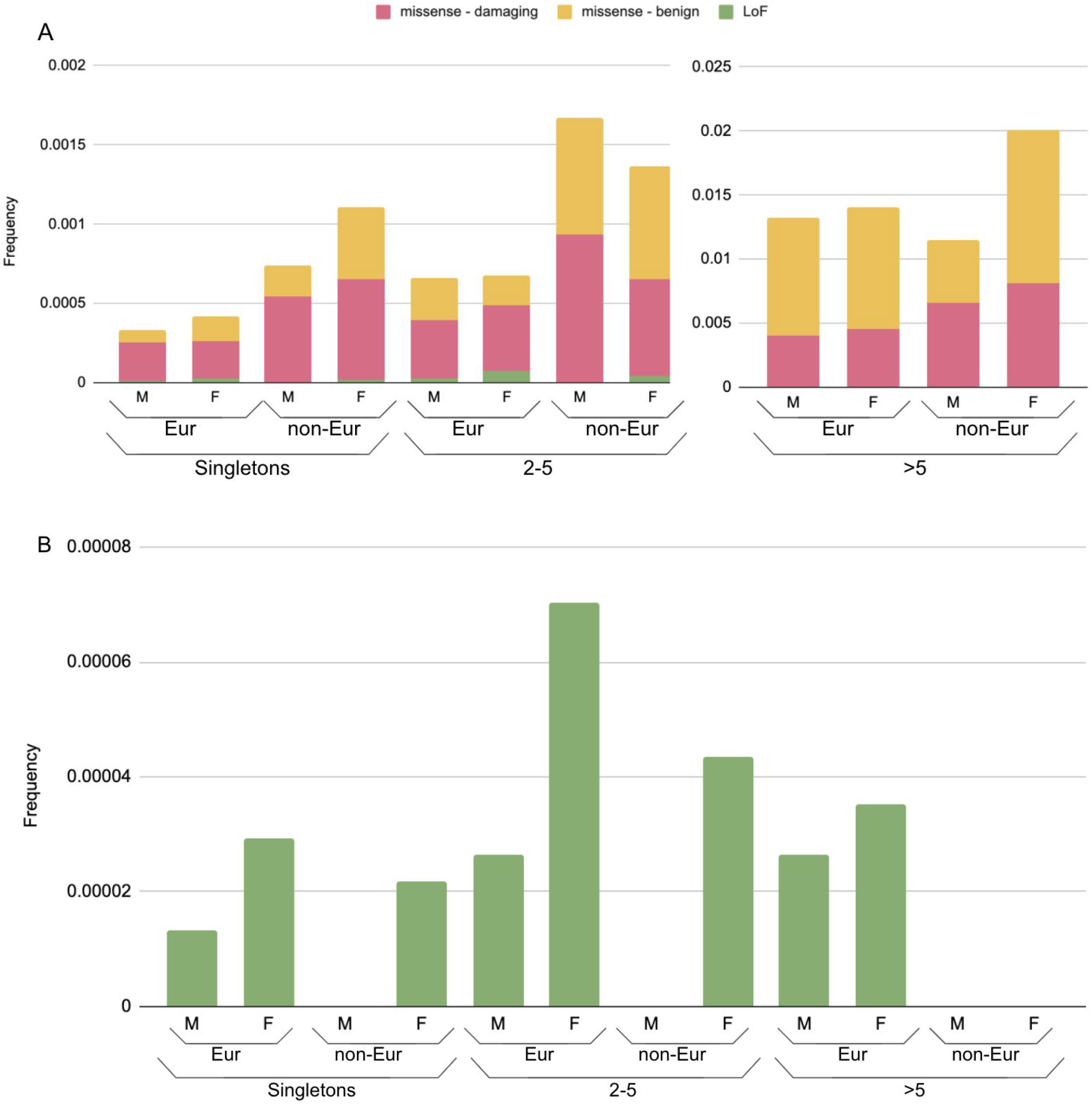
Aggregated frequency of variants found in the Helix dataset of >200,000 unrelated individuals. A) Proportion of individuals in Helix dataset of European and other (non-Eur) ancestries carrying coding variants; (B) carrying only LoF variants. Frequencies are broken up according to the number of carriers seen in the Helix dataset (singletons, 2-5 carriers, or >5 carriers). M=male, F=female.

**Figure 4.**
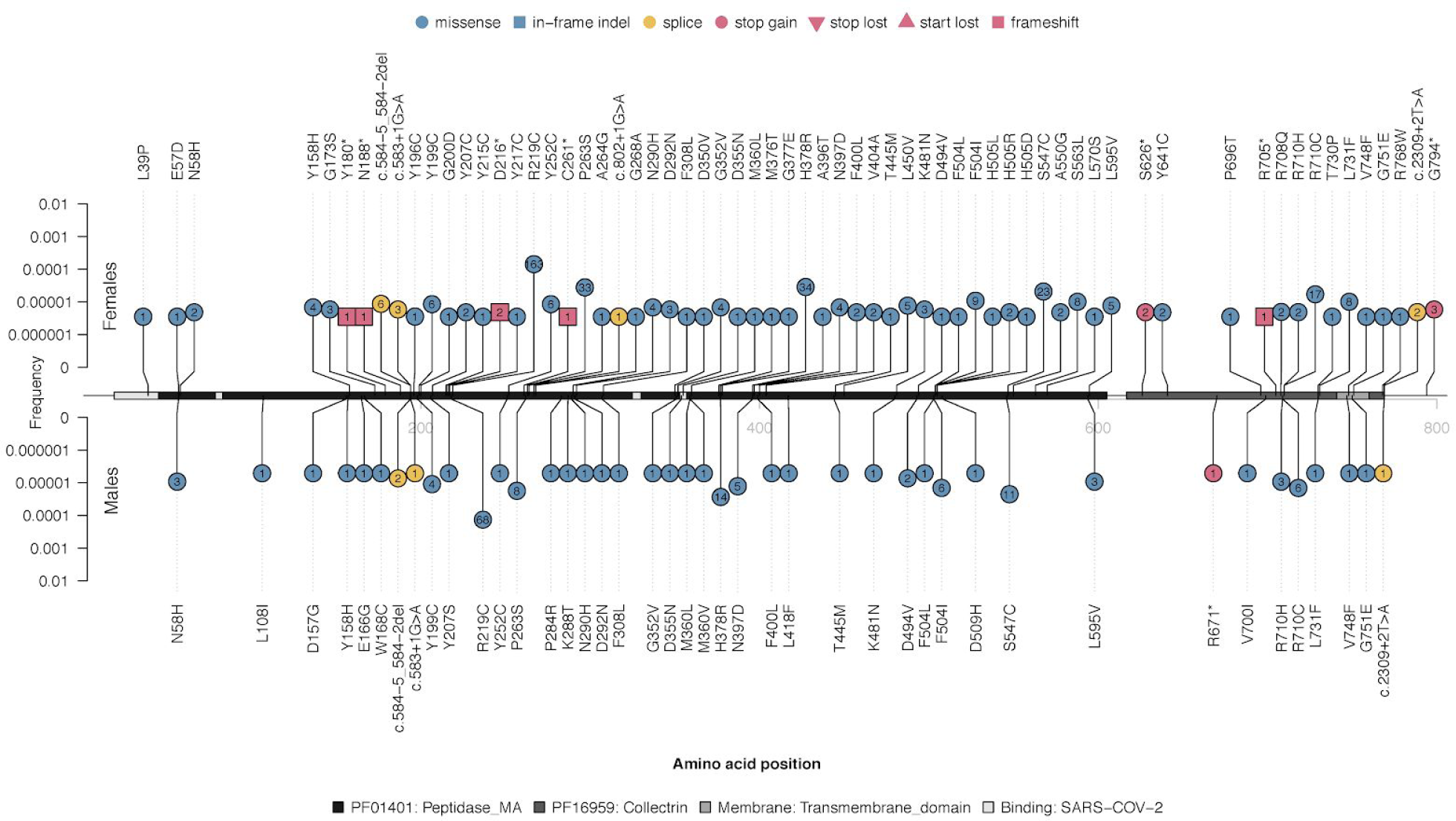
Lolliplot^18^ of damaging coding variants found in *ACE2*. Variants shown are from the unrelated European ancestry individuals in the Helix dataset. The amino acid positions along ACE2 are shown at the horizontal middle line together with the different PFAM domains^26^, the C-terminal transmembrane domain, and the SARS-CoV-2 binding regions^16^ shown as gray boxes. Vertical lines represent variant frequencies (log scale) in females (above) and males (below). The type of variant is indicated by the shape and color of each lollipop, as well as the number of carriers within each. Variants were defined as damaging according to a consensus between SIFT and Polyphen2 (as in Figure 2)^19–22^. Y axis is log scale.

## *ACE2* CNVs

Of the CNVs identified in our Helix dataset, besides the whole-gene deletion in one individual noted above, we found two individuals who also had a single exon deleted each and 0.02% of our population who had a duplication of the first 6-7 exons of the gene, each of which also has the potential to cause loss of function. All of these CNVs were found in females. The only CNV that we found in males was a duplication of the entire *ACE2* gene, which is duplicated in ~0.04% of our population. This is consistent with the frequency of this duplication shown in gnomAD^15^. As a whole-gene duplication could increase expression, we compared the genotypes in our dataset to those in the GTEx database^11^. In GTEx, there is a very common haplotype associated with differential expression of the gene in some tissues, extending from the 5’-end of the gene to far upstream and into the next gene, an *ACE2* homolog with 45% identity to *ACE2* but no catalytic domain^27^. We found that in the Helix dataset, the carriers of the rare whole-gene duplication were on the very common low-expression *ACE2* haplotype. Although this does not mean that duplication carriers have low expression of the gene, it does show that the carriers of this rare duplication variant are not the drivers of the association between expression and that haplotype, as does the spread-out nature of the common variant’s violin plot (Figure 5).

**Figure 5.**
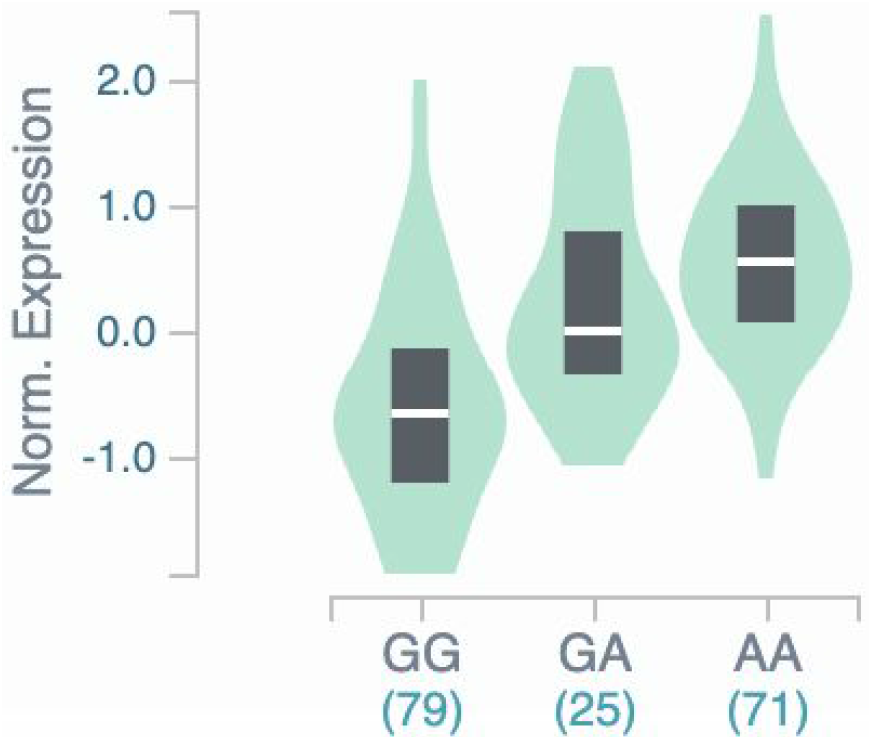
Association between ACE2 variant rs4830974 and Brain Frontal Cortex (BA9) gene expression. Exemplar genotype for low-expression eQTL from GTEx^11^, p=2.6×10-17. All other eQTL in LD with this genotype had similar expression levels.

## PheWAS

Finally, we looked into non-COVID-19-related phenotypes that might be associated with variation in *ACE2*. In our recent analysis of rare coding variants in the Healthy Nevada Project and UK Biobank ^28–30^, we did not identify any genome-wide significant associations between the >4,000 phenotypes analyzed and the collapsed rare coding variants in *ACE2*. However, we highlight here the top 20 associations from the meta-analysis across the two cohorts (Table 1).

**Table 1.**
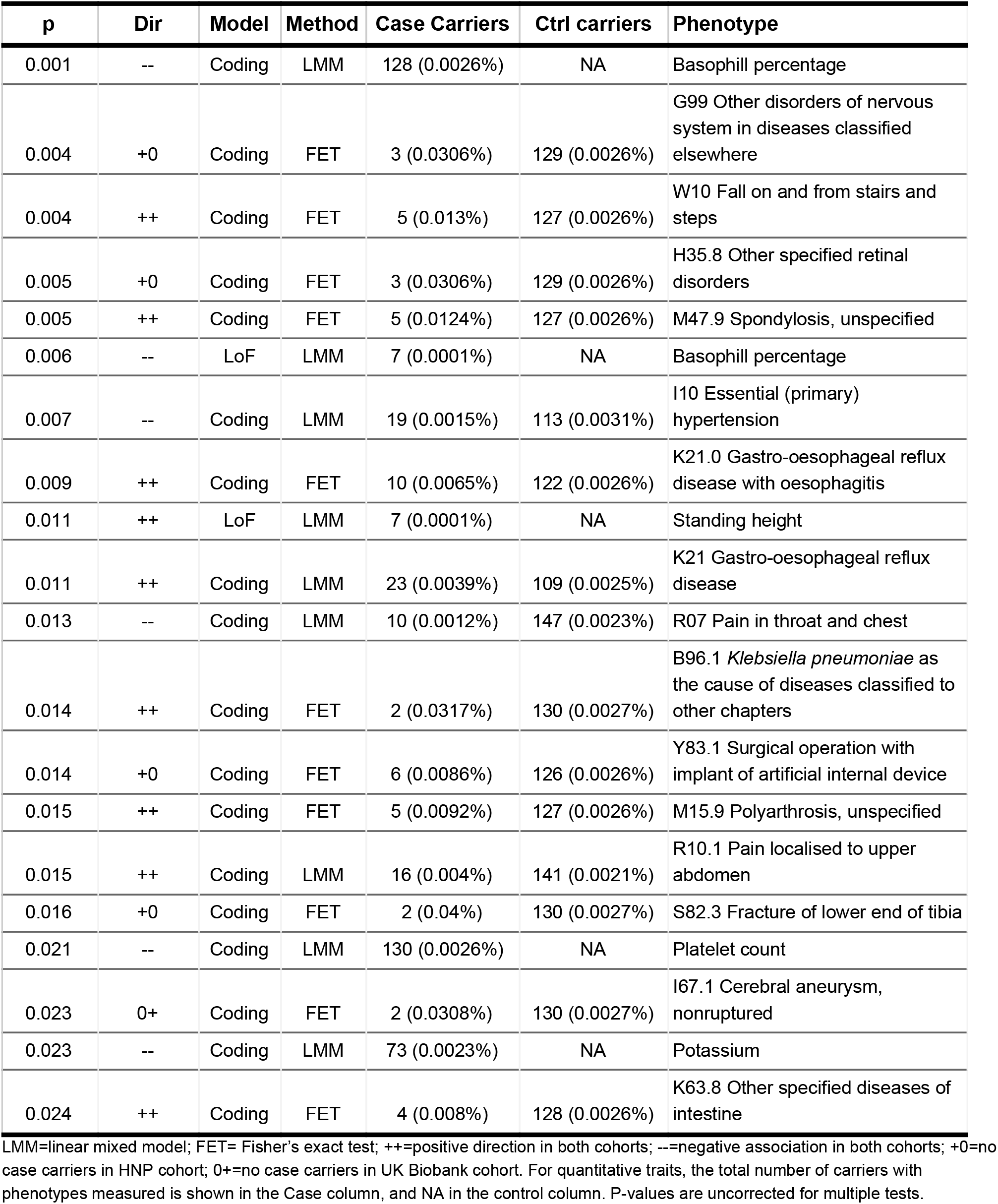
Top 20 phenotypes associated with *ACE2* in UKB and HNP

Consistent signals were seen in both cohorts for a weak association between those carrying coding variants in *ACE2* and having lower basophil percentages. Some of the other signals observed included a weak protective effect of coding variants for hypertension, which is interesting given this gene’s function in blood pressure and the known relationship between the related gene *ACE* and human blood pressure^31^; and a predisposition for those with coding variants to have gastro-oesophageal reflux disease with oesophagitis, which is consistent with the nominal association observed between the low-frequency intronic variant rs183546232 in *ACE2* (MAF 1%) and eosinophilic esophagitis in an independent dataset^32^. There was also a nominal association between coding variants and ICD10 code B96.1, *Klebsiella pneumoniae* as the cause of diseases classified elsewhere, but this was driven by only 2 case carriers across both cohorts (although Fisher’s exact test is robust to small samples). Investigations of additional pneumonia-related ICD10 codes such as J18 revealed no significant association with *ACE2* variation.

We also utilized the existing publicly available UK Biobank GWAS results from the Neale lab to perform a pheWAS of common variants in *ACE2^33^*. We did not identify individual common variants in this gene with significant associations with any phenotypes. We anticipate, with the sizable global scientific efforts dedicated to COVID-19 host genomics^2,3^, that potential associations between *ACE2* variants and COVID-19 response will be characterized once human genetic data paired with COVID-19 phenotypes are generated.

## Summary

In conclusion, there is moderate evidence that *ACE2* variation is associated with gastro-oesophageal reflux disease with oesophagitis, and limited evidence of it having a strong role in other phenotypes in the absence of COVID-19. Homo/hemizygous LoF of *ACE2* appears to be infrequently tolerated and is likely extremely deleterious. Although LoF variants in *ACE2* are extremely rare, variants affecting the regions that interact with SARS-CoV-2 are more prevalent, and the impending research studies on human genetics and COVID-19 response may identify some of these variants as conferring resistance or heightened susceptibility to the virus.

## Acknowledgments

We acknowledge the contributions of Y. Ni, S. Bobulsky, S. White, M. Isaksson, F. Tanudjaja, Helix participants, and the entire Helix laboratory and bioinformatics teams.

## Notes

https://s3.amazonaws.com/helix-research-public/data_releases/ace2_variants_helix_db.txt

### Competing Interest Statement

All authors are employees of Helix.

### Summary of Updates

Figure 4 revised, allele frequency corrected

